# Combination of loop-mediated isothermal amplification and AuNP-oligoprobe colourimetric assay for pork authentication in processed-meat products

**DOI:** 10.1101/2020.07.12.199091

**Authors:** Pattanapong Thangsunan, Sasithon Temisak, Phattaraporn Morris, Leonardo Rios-Solis, Nuttee Suree

## Abstract

Pork adulteration is a major concern for Muslims and Jews whose diets are restricted by religious beliefs, as well as those who are allergic to pork meat and its derivatives. Accurate pork authentication is of great importance to assist this demographic group of people in making decision on their product purchase. The aim of this study was to develop a new analytical method for pork authentication in processed-meat products based on a combination of loop-mediated isothermal amplification (LAMP) and AuNP-nanoprobe colourimetric assay. The LAMP conditions were first optimised to obtain the highest yield of amplified DNA products within the shortest time. Oligoprobe-functionalised AuNPs were then hybridised with LAMP-DNA amplicons, and subsequently challenged with MgSO_4_ at a high concentration to induce AuNP aggregation. In the presence of pork DNA, the colloidal AuNPs-probe remained unchanged in its red colour, which indicates the dispersion of AuNPs. In contrast, in the absence of pork DNA, the colour was changed to colourless as a result from the aggregation of AuNPs. The LAMP-AuNP-nanoprobe assay offers a high sensitivity with a limit of detection as low as 100 pg of pork DNA. The assay is highly specific to pork content without cross-reactivity with the other meat species tested. The assay developed herein can become a simple, inexpensive, precise, and rapid analytical tool for small laboratories or the general public interested in halal food authentication.

## Introduction

Consumers have generally been concerned about the composition of food they consume, particularly due to the preference on meat products, religious beliefs, as well as health issues (Alikord et al. 2017; Ballin 2010; Farag et al. 2015). Over the past decade, adulteration in meat products has also been widely reported (Kane and Hellberg 2016; Pinto et al. 2015). Meat adulteration can include substitution of higher-valued meats with lower-valued ones, intentional mixing of undeclared meat types or ingredients, additions of plant proteins instead of meat content, and unintentional contamination during processing (Barakat et al. 2014; Cawthorn et al. 2013). These activities may negatively affect consumers’ trust on food quality and safety. Therefore, accurate labelling of food products will not only help to protect the consumers from fraudulent actions, but also promote fair trades in local and international levels (Alikord et al. 2017; Barakat et al. 2014; Nešić et al. 2017). For this reason, approaches for identification of meat species has been of great interest in order to support the food industry and to help customers in making decision on food products.

Many analytical techniques, based on anatomical, histological, microscopic, organoleptic, electrophoretic, chromatographic, and immunological principles have been developed for the authentication of meat species (Nešić et al. 2017). However, many of these techniques exhibit low sensitivity or are unsuitable for complex and processed food samples (Alikord et al. 2017). DNA detection methods have been of great interest since the structure of DNA is highly stable at high temperature, especially during the food processing steps, while proteins can be denatured, rendering them unsuitable analytes for accurate identification (Ballin et al. 2009). The most frequently used DNA detection techniques for meat identification are based on polymerase chain reaction (PCR) such as randomly amplified polymorphic DNA – PCR (RAPD-PCR), restriction fragment length polymorphism - PCR (RFLP-PCR), PCR with species-specific primers, real-time PCR/quantitative PCR (RT-PCR/qPCR) and digital PCR (dPCR) (Alikord et al. 2017; Ballin et al. 2009; Farag et al. 2015). Despite their high sensitivity and specificity, PCR methods have some limitations including their complicated experimentation, the need of a thermal cycler or an additional detection machine, their relatively high cost of reagents and time consumption, and the need of skilled operators (Ali et al. 2012; Ckumdee et al. 2016; Seetang-Nun et al. 2013; Takano et al. 2017; Tomita et al. 2008). This thus presents an opportunity for developing novel and more efficient approaches for authenticating meat species.

In 2000, Notomi and colleagues developed a novel nucleic acid amplification technique called the loop-mediated isothermal amplification (LAMP) (Notomi et al. 2000). LAMP can efficiently work at a constant temperature using a DNA polymerase enzyme with strand displacement ability together with a set of four primers designed from six distinct regions on the target DNA sequence (Notomi et al. 2000; Tomita et al. 2008). The technique is known for its high specificity and selectivity, high cost effectiveness, fast processing time, and independency from expensive equipment (Tomita et al. 2008). As a result, LAMP has been used in a wide range of applications to overcome the limitations of PCR (Becherer et al. 2020; Kundapur and Nema 2016; Martzy et al. 2019; Mori et al. 2013; Tasrip et al. 2019). Among those applications, LAMP can be combined with other systems to allow better detection of the amplified products. For example, gold nanoparticle (AuNP) colourimetric assay is a visual detection platform based upon aggregation and dispersion of gold nanoparticles (Aldewachi et al. 2018). AuNPs possess a unique property known as surface plasmon resonance (SPR). The increase of AuNP size induces the interparticle plasmon coupling, causing a red shift of the SPR band towards a longer wavelength (Aldewachi et al. 2018; Radwan and Azzazy 2009; Zeng et al. 2011; Zhao et al. 2008). Therefore, as the formation of AuNP aggregates occurs, a transition of solution colour from red to blue or purple can be visible to the naked eyes (Aldewachi et al. 2018; Zhao et al. 2008). Stability of AuNPs in a solution can be modulated through crosslinking and non-crosslinking mechanisms (Zhao et al. 2008). Biological events such as an interaction of antigen-antibody and a hybridisation of DNA can be used to stabilise or destabilise colloidal AuNPs by controlling the net potential between interparticle attractive and repulsive forces (Aldewachi et al. 2018; Glomm 2005; Napper 1984; Vasconcelos et al. 2005; Walker and Grant 1996; Zhao et al. 2008). While the LAMP method generates DNA amplicons, its combination with the AuNP colourimetry can enable a clear and sensitive detection of DNA or RNA of interest. To this end, the LAMP - gold nanoprobe methods have been applied successfully for a wide range of fields including diagnosis of diseases (Lou et al. 2020; Najian et al. 2016; Qiuhua et al. 2014), microbial drug resistance (Ckumdee et al. 2016; Veigas et al. 2013), forensic sciences (Watthanapanpituck et al. 2014), and pathogen detection in foods (Garrido-Maestu et al. 2017; Kong et al. 2018; Seetang-Nun et al. 2013; Suebsing et al. 2013; Wachiralurpan et al. 2018).

Pork, aside from being one of the most widely consumed meats in the world (FAO 2019; USDA 2018), tends to be a source of adulteration in other meat products such as beef and lamb, mainly due to its cheaper price and similarity in colour and texture (Barakat et al. 2014; Wissiack et al. 2003). Accurate pork authentication is not only necessary for Muslims and Jews, whose religious beliefs highly influence the choice of meat consumption (Barakat et al. 2014; Sheikha et al. 2017), but also for those who are allergic to pork and pig-derived product varieties (Fournier et al. 2011; Mamikoglu 2005). In this study, a new analytical platform based on a combination of loop-mediated isothermal amplification with AuNP-nanoprobes was developed in order to authenticate pork content in commercial processed-meat products. With its high sensitivity, specificity, efficiency, and simplicity, requiring basic instruments like water baths and heating blocks, this novel method has high potential to become widely used both in food laboratories and in the food industry.

## Materials and methods

### Sample preparation and DNA extraction

Fresh pork (*Sus scrofa*), chicken (*Gallus gallus*), duck (*Anas platyrhynchos*), sheep (*Ovis aries*), salmon (*Salmo salar*) and other processed meat products (2 samples of pork sausage, pork ball, chicken burger, beef burger and shrimp dumpling) were purchased from supermarkets in Pathum Thani, Thailand. Fresh beef meat (*Bos taurus*) was obtained from a halal food market in Pathum Thani, Thailand. All meat samples were individually blended and freeze-dried (CHRIST, Gamma 1-16 LSC, Germany). The dried meat samples were then sieved through 300 μm standard test sieves (Retsch, Fisher Scientific, USA) and stored at −80 °C until further use.

To obtain genomic DNA, 2 g of meat or processed food powder was extracted and purified using DNeasy Mericon Food Kit (Qiagen, Germany) according to the manufacturer’s instructions. After that, purified DNA samples were determined for their concentration and purity using UV-Vis spectroscopy with a microplate reader (Tecan Spark, Switzerland).

### Plasmid construction

The partial mitochondrial DNA sequences of *Sus scrofa* (GenBank accession no. AF034253) and *Bos taurus* (GenBank accession no. AY526085) were amplified using the primers shown in Table 1. The expected size of PCR products for *S. scrofa* and *B. taurus* was 622 bp and 585 bp, respectively. The PCR reaction with a total volume of 25 μl was prepared as follows: 1x of PCR buffer, 0.2 μM dNTP mix, 1.5 mM MgCl_2_, 0.3 μM each of forward and reverse primers (Table 1), 50 ng of pork or beef genomic DNA and 2.5 U *Taq* DNA polymerase (Invitrogen, USA). PCR cycle started with initial denaturation at 94 °C for 3 min, followed by 35 cycles of denaturation at 94 °C for 45 s, annealing at 55 °C for 30 s and extension at 72 °C for 45 s, which then ended with a final extension at 72 °C for 10 min. The PCR products were analysed using 1.5% agarose gel electrophoresis in 1x Tris-borate-EDTA (TBE) buffer and purified from the gel using a NucleoSpin^®^ gel and PCR clean-up kit (Macherey-Nagel, Germany).

**Table 1.**
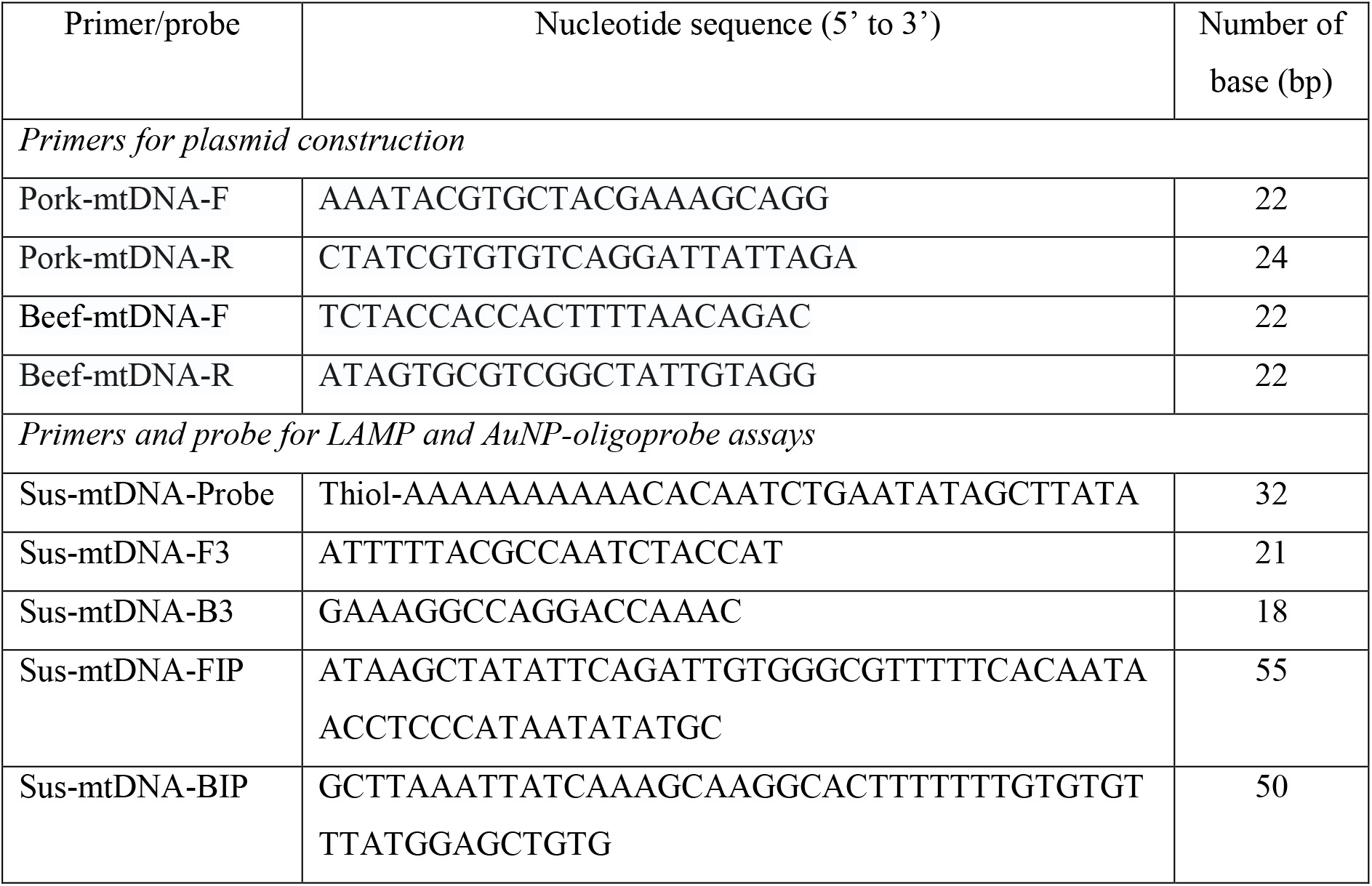
Nucleotide sequences for the primers and oligoprobe used in this study.

To construct plasmids, the PCR fragments were ligated into pCR^®^ 2.1 vector (Invitrogen, USA) according to the manufacturer’s instructions. The ligated plasmids were transformed into One Shot™ Top10F’ Chemically Competent *E. coli* (Invitrogen, USA) by heat-shock method. The *E. coli* was grown on LB agar plus 100 μg/ml ampicillin prior to blue-white colony selection. Some of the white colonies were picked and cultured in LB broth plus 100 μg/ml ampicillin. The plasmids were purified using a QIAprep^®^ Spin Miniprep kit (Qiagen, Germany) according to the manufacturer’s protocol. Purified plasmid concentration was examined using a NanoDrop One UV-Vis spectrophotometer (Thermo Scientific, USA).

### LAMP primer design

The partial mitochondrial DNA sequence of *S. scrofa* was exploited to design a set of LAMP primers via Primer Explorer version 5 (https://primerexplorer.jp/e/v5_manual/index.html). The details of primer locations and sequences are presented in Fig. S1 and Table 1. All of the primers used were synthesised and purified using PAGE by Macrogen Inc., (Korea).

### LAMP assay

The method for LAMP was modified from the previously published work (Tomita et al. 2008). LAMP reactions at a final volume of 12.5 μl were prepared. Each reaction contained 1.6 μM each of the inner primers (SUS-mtDNA-FIP and SUS-mtDNA-BIP in Table 1), 0.2 μM each of the outer primers (SUS-mtDNA-F3 and SUS-mtDNA-B3 in Table 1), 1.4 mM of dNTP mix, 6 mM of MgSO_4_, 0.4 M of Betaine (Sigma-Aldrich, USA), 8 U of *Bst* DNA polymerase (large fragment; New England Biolabs Inc., Beverly, MA, USA), 1x of the supplied buffer and a variable amount of DNA template. Plasmid DNA containing *S. scrofa* mitochondrial DNA sequence (pCR-pork-mtDNA plasmid – Fig. S2) was used as a positive control while sterile ddH_2_O was utilised as a no-template control (NTC) for the entire experiments. All of the LAMP reactions were performed at 61°C for 45 min unless stated otherwise, followed by enzyme inactivation at 90 °C for 2 min. All of the experiments were replicated in at least triplicate. The LAMP products were analysed using 2% agarose gel electrophoresis in 1x TBE buffer.

### Functionalisation of AuNPs

The sequence used for designing a ssDNA probe was obtained from mitochondrial DNA of *S. scrofa* and thiolated at the 5’-end (Table 1). 20 nm AuNPs (Sigma-Aldrich, USA) were functionalised with the thiolated oligoprobe according to the method modified from the previously published work (Hill and Mirkin 2006). In brief, 1 ml of 1 nM of the colloidal AuNPs solution was incubated with 4 nmol of freshly DTT-reduced oligoprobe overnight at room temperature with gentle agitation. Phosphate buffered saline (PBS, pH 7.0) and sodium dodecyl sulfate (SDS) were sequentially added to the AuNPs-oligoprobe solution to a final concentration of 9 mM and 0.1% (w/v) respectively. Salting buffer containing 10 mM PBS and 2 M NaCl was then added to the solution by making six additions over the course of two days to reach a final NaCl concentration of 0.3 M. After the last NaCl addition and overnight incubation, the oligoprobe-coated AuNPs were washed three times in 500 μl of assay buffer containing 10 mM PBS, 150 mM NaCl, 0.1% (w/v) SDS, pH 7.4. The functionalised AuNPs were resuspended in the assay buffer to obtain a stock concentration of 2 nM and stored in the dark at room temperature until further use.

### AuNPs-oligoprobe colourimetric assay for a proof of concept

The proof of concept experiment was conducted in a reaction with a total volume of 10 μl. In each reaction, 5 μl of AuNPs-nanoprobe was incubated with 2.5 μl of 10 μM ssDNA target at 61 °C for 15 min. The sequence for ssDNA complementary to the nanoprobe was 5’-TATAAGCTATATTCAGATTGTG-3’, whereas that for non-complementary ssDNA was 5’-AACTCGGATACCCAAGGTCTTAC-3’. No-template control (NTC) for this experiment was sterile ddH_2_O. After hybridisation, the solution was mixed with 0.5 M MgSO_4_ and incubated for 30 min at room temperature. The transition of AuNPs-nanoprobe solution colour was visually observed and the absorbance of 400-700 nm was examined using a microplate reader (Tecan Spark, Switzerland). The experiments were replicated in at least triplicate.

### AuNPs-oligoprobe colourimetric assay for LAMP product testing

The colourimetric assays were performed in a total reaction volume of 10 μl. In each reaction, 5 μl of oligoprobe-functionalised AuNPs was incubated with 2.5 μl of LAMP products at 61 °C for 15 min. After hybridisation, 2.5 μl of 2 M MgSO_4_ was added and the solution was incubated for 30 min at room temperature. The change of solution colour was determined by visual observation and UV-Vis spectroscopy between 400 and 700 nm using a Spark^®^ multimode microplate reader (Tecan, Switzerland). All of the experiments were replicated in at least triplicate.

### Primer-specific PCR for meat-species specificity

The primers for specific PCR were obtained from the previous study (Montiel-Sosa et al. 2000), which were designed for pork D-loop mitochondrial DNA. The sequence for the forward primer (Pig-F) was 5’-AACCCTATGTACGTCGTGCAT -3’, whereas that for the reverse primer (Pig-R) was 5’-ACCATTGACTGAATAGCACCT -3’. Each 25 μl reaction of PCR contained 1x *Pfx* amplification buffer, 1 mM MgSO_4_, 0.3 mM dNTP mix, 0.3 μM each of forward and reverse primers, 20 ng DNA template and 1.25 U Platinum *Pfx* DNA polymerase (Invitrogen, USA). The PCR cycle started from 94 °C for 2 min, followed by 30 cycles of 94 °C for 15 s, 58 °C for 30 s and 68 °C for 45 s. The PCR was ended with final extension at 68 °C for 5 min. The PCR products were analysed with 1.5% agarose gel electrophoresis in 1x TBE buffer.

### Real-time PCR for meat-species specificity

The primers and probe for RT-PCR were used according to the previous work (Köppel et al. 2011), which were specific for pork beta-actin gene (GenBank accession no. DQ452569). The sequence for pork_F primer (forward direction) was 5’-GGAGTGTGTATCCCGTAGGTG-3’ while that for pork_R primer (reverse direction) was 5’-CTGGGGACATGCAGAGAGTG-3’. The sequence for the probe was 5’-FAM-TCTGACGTGACTCCCCGACCTGG-BHQ1-3’. In a total volume of 20 μl, each reaction contained 1x TaqMan Universal PCR mastermix (Applied Biosystems by Thermo Fisher Scientific, USA), 300 nM each of forward and reverse primer, 200 nM of probe, and 30 ng of DNA template. The RT-PCR was performed using a thermo cycler (ABI 7500 Real-Time PCR, Thermo Fisher Scientific, USA) at 95 °C for 10 min, followed by 45 cycles of 95 °C for 15 s and 58 °C for 1 min. The RT-PCR data were analysed using 7500 Software v2.3 (ABI 7500 Real-Time PCR).

## Results and discussion

### Optimisation of condition for LAMP assays

To achieve the highest performance of LAMP assays, two parameters for LAMP, melting temperature (T_m_) and incubation period, were optimised. To examine the optimal T_m_, during performing LAMP reactions, the temperature was varied between 59 °C and 65 °C. When the plasmid containing pork mitochondrial DNA (pCR-pork-mtDNA plasmid – Fig. S2) was used as a template, LAMP products from the reactions at 59, 61 and 63 °C showed a similar pattern with intense bands between 200-400 bp (Fig. S3). However, the LAMP products obtained from the 65 °C reaction showed a slightly different pattern with faint 200-400 bp bands, indicating less yield of LAMP products (Fig. S3). No band was observed when using the plasmid carrying beef mitochondrial DNA sequence (pCR-beef-mtDNA plasmid – Fig. S4) as a negative control template (Fig. S3). Since *Bst* DNA polymerase has the optimal temperature between 60 and 65 °C (Notomi et al. 2000), the T_m_ of 61 °C was selected as an optimal condition for LAMP assays.

The incubation period was also studied as one of the parameters for LAMP. The LAMP T_m_ was fixed at 61 °C with incubation period varied from 15 to 60 min. Using pCR-pork-mtDNA plasmid (Fig. S2) as a template, LAMP products were detected at 30 min and the amount of the amplified products increased as the reaction time increased (Fig. S5). The yield of the LAMP products obtained seemed to reach the highest point at 45 min of incubation (Fig. S5). The reactions with pCR-beef-mtDNA plasmid (negative control) and ddH_2_O (no template control) did not show any LAMP amplicons (Fig. S5). The optimal incubation time for LAMP in this study was 45 min. Conclusively, the LAMP assays were performed at 61 °C for 45 min.

### Proof of concept for AuNP colourimetric assays

A proof of concept of AuNP colourimetric assays was performed to assure its detectability in the presence of the target and non-target DNA. AuNPs (20 nm diameter) were coated with a ssDNA probe with a nucleotide sequence specific to pork mitochondrial DNA (Fig. S1). A thiol group was introduced to the 5’-end of the probe to allow the oriented conjugation of the probe on the surface of AuNPs via thiol-metal interactions (Weisbecker et al. 1996).

Functionalised AuNPs were then hybridised with the ssDNA target. After hybridisation, MgSO_4_ was added as a salt to challenge the stability of AuNPs upon the change of ionic strength of the solution (Burns et al. 2006; Ckumdee et al. 2016; Seetang-Nun et al. 2013). In the presence of ssDNA target with the complementary sequence to the oligoprobe, the solution colour remained red as its original (Fig. 1A), indicating the dispersion of AuNPs. However, with the non-complementary ssDNA target (negative control) or in the absence of ssDNA (no template control), the solution changed to colourless (Fig. 1A) as a result from the aggregation of AuNPs. UV-Vis spectroscopy was also used to monitor the behaviour of functionalised AuNPs in the presence or absence of the target (Fig. 1B). Surface plasmon spectrum of the AuNPs-probe with a complementary target showed a distinct peak at 520 nm (Fig. 1B), which is common for colloidal 20 nm AuNPs (Carrillo-Cazares et al. 2017; Haiss et al. 2007). On the contrary, in the absence of a specific target, this spectrum was shifted towards a longer wavelength (Fig. 1B). This event can be explained by the fact that the binding of complementary ssDNA target to the probe on AuNP surface increases electrostatic repulsion forces and steric hindrance from negative charges on phosphate backbone of DNA, leading to high stability of AuNPs-probe even though being in high salt concentration environment (Aldewachi et al. 2018; Seetang-Nun et al. 2013). Therefore, the AuNPs solution remained red in colour. Without ssDNA target, the charges of AuNPs-probe are intolerant to the increase of ionic strength, resulting in AuNPs aggregation. Aggregation of AuNPs can be visually detected by the change of solution colour from red to blue/purple (Aldewachi et al. 2018; Pamies et al. 2014; Seetang-Nun et al. 2013). In some circumstances, AuNPs colour can also turn colourless if the aggregation is extreme, causing the shift of SPR bands to the infra-red region (Aldewachi et al. 2018).

**Fig. 1.**
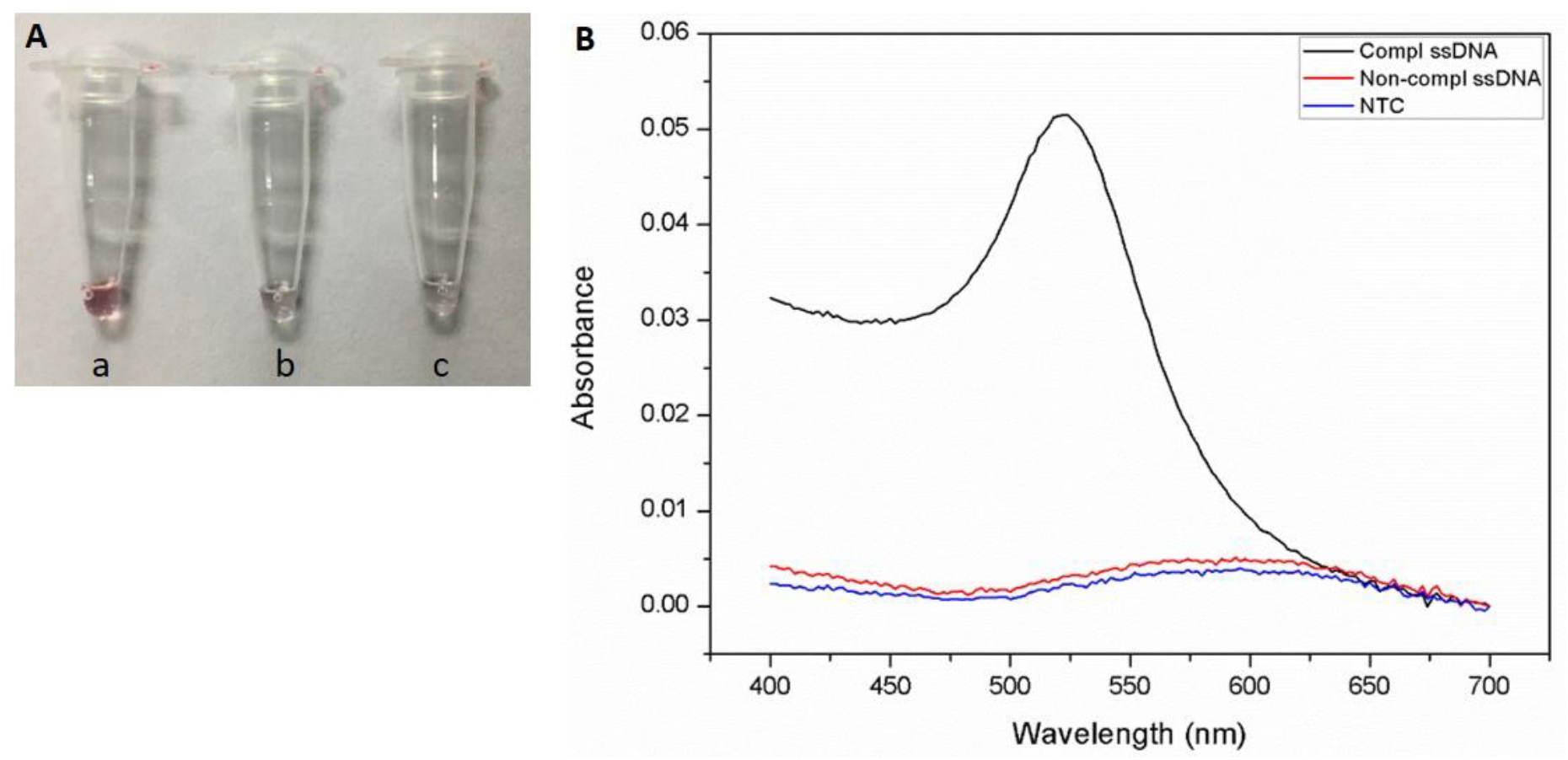
The proof of concept for AuNP colourimetric assays. (A) Digital photograph showing the change of colloidal AuNP colour after adding 0.5 M MgSO_4_ to the solution with (a), 10 μM of complementary ssDNA; (b), 10 μM of non-complementary ssDNA and (c), sterile ddH_2_O as a no-template control. (B) UV-Vis spectra of AuNP-probe with complementary ssDNA, non-complementary ssDNA and no-template control.

### Combination of LAMP and AuNP-probe colourimetric assays

In this study, LAMP as a tool for target gene amplification, and AuNP-probe colourimetric assay as a method for visually detecting the presence of a target of interest, were combined to improve the efficiency of pork DNA detection. Two parameters affecting the performance of the assays, which were MgSO_4_ concentrations and hybridisation temperature, were optimised.

### Effects of MgSO_4_ concentrations

The effects of MgSO_4_ concentrations on the aggregation of AuNPs-probe were studied since the ionic strength of salt in aqueous media can modulate the stability of colloidal AuNPs as observed by the transition in colour of colloidal AuNPs (Burns et al. 2006; Pamies et al. 2014; Seetang-Nun et al. 2013). In the present work, MgSO_4_ was chosen as a salt to induce AuNP aggregation. After hybridisation, the solution of MgSO_4_ at a concentration of 0.25, 0.5, 1 and 2 M was added to the mixture of AuNPs-probe and LAMP amplicons. As shown in Fig. 2A, at 0.25 and 0.5 M MgSO_4_, AuNPs-probe colour remained red no matter the DNA templates for LAMP were pCR-pork-mtDNA plasmid (positive control), pork DNA, beef DNA (negative control) or ddH_2_O (NTC). Using 1 M MgSO_4_, a slight drop of AuNPs-probe colour from red to pale red was detected for beef gDNA and NTC reactions while the solution colour for pCR-pork-mtDNA plasmid and pork gDNA remained unchanged (Fig. 2A). A significant change of AuNPs colour was observed at 2 M MgSO_4_ where the solution colour was still red for LAMP products from pCR-pork-mtDNA plasmid and pork gDNA, but became colourless for beef DNA and NTC (Fig. 2A). These data were in good agreement with the UV-Vis spectroscopic data (Fig. 2B). The A520/A650 ratios for AuNPs-probe with beef DNA and NTC were decreased with 1 and 2 M MgSO_4_ whereas those for pCR-pork-mtDNA plasmid and pork DNA remained unchanged (Fig. 2B). No significant changes of the A520/A650 ratio for all four samples were observed for 0.25 or 0.5 M MgSO_4_ (Fig. 2B). Thus, 2 M MgSO_4_ was selected as the optimal concentration of salt used for inducing AuNP aggregation.

**Fig. 2.**
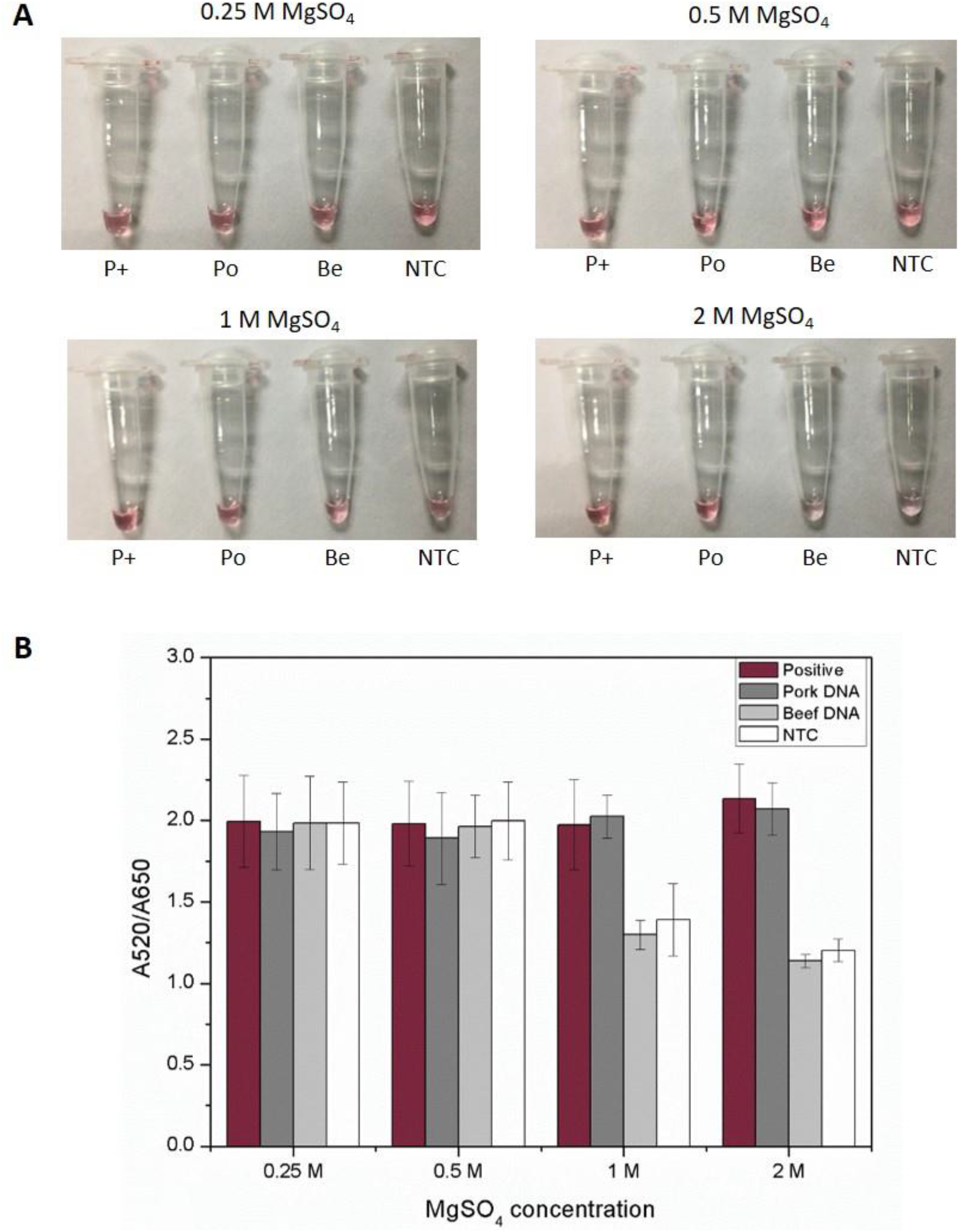
Effects of MgSO_4_ concentrations on AuNPs-probe aggregation. (A) Digital photographs presenting AuNPs-probe after mixing with different concentrations of MgSO_4_. LAMP products to be tested were amplified from (P+), 25 ng of pCR-pork-mtDNA plasmid as a positive control; (Po), 40 ng of pork genomic DNA; (Be), 40 ng of beef genomic DNA and (NTC), sterile ddH_2_O as a no-template control. (B) A520/A620 ratio of AuNPs-probe after adding different concentrations of MgSO_4_. Data are means ± SD (n = 8).

### Effects of hybridisation temperature

Hybridisation temperature is one of the key factors to achieve AuNPs-probe colourimetric assays since it governs a specific interaction between two complementary strands of DNA (Ge et al. 2012; Harris and Kiang 2006; Hottin et al. 2009). In this study, the mixtures of LAMP amplicons and AuNPs-probe were hybridised at room temperature, 50 and 61 °C. Fig. 3A revealed that at hybridisation temperature of 50 and 61°C, the aggregation of AuNPs was observed in LAMP reactions from beef DNA and NTC while no aggregates were seen with LAMP products from pCR-pork-mtDNA plasmid (positive control) and pork DNA. However, the rate of AuNP aggregation at 61 °C was faster than that of 50 °C. At room temperature, a slow AuNPs aggregation was found for AuNPs with beef DNA and NTC as observed by a slight drop of red colour of AuNPs (Fig. 3A). This is likely because the rate of hybridisation of LAMP amplicons to oligoprobe takes place more rapidly at higher temperature. The UV-Vis spectroscopic measurement agreed with visual observation results that the temperature of 61 °C was optimum since it caused the most significant drop of the A520/A650 ratio for non-pork DNA samples (Fig. 3B) whereas the ratios of A520/A650 for pCR-pork-mtDNA plasmid and pork DNA remained unchanged for all the hybridisation temperatures tested.

**Fig. 3.**
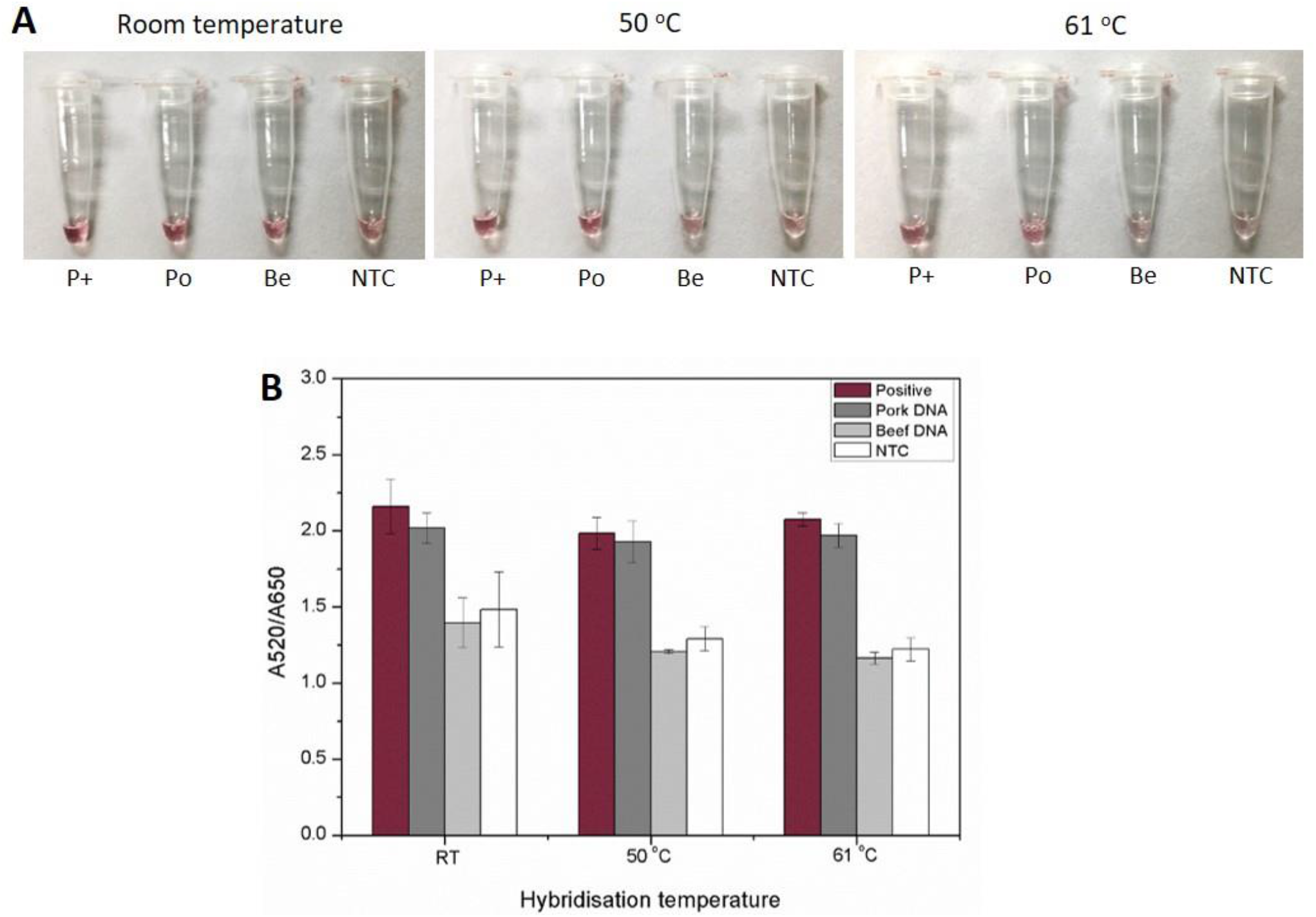
Effects of hybridisation temperature on AuNPs-probe aggregation. (A) Digital photographs showing AuNPs-probe solution colour after hybridisation at room temperature, 50 and 61 °C. LAMP products to be tested were amplified from (P+), 25 ng of pCR-pork-mtDNA plasmid as a positive control; (Po), 40 ng of pork genomic DNA; (Be), 40 ng of beef genomic DNA and (NTC), ddH_2_O as a no-template control. (B) A520/A620 ratio of AuNPs-probe after hybridisation at different temperatures. Data are means ± SD (n = 6).

### Specificity of LAMP-AuNPs colourimetric assays

Specificity of LAMP-AuNP-probe assays was also investigated since cross-reactivity is likely to cause false positives in non-pork containing samples. LAMP reactions were performed with genomic DNA from pork, beef, chicken, duck, sheep and salmon. The results from agarose gel electrophoresis showed that LAMP products were only seen in the reactions with pCR-pork-mtDNA plasmid (positive control) and pork DNA while there were no amplified LAMP products in the non-pork DNA samples (Fig. 4A). The LAMP products were then tested using AuNP-probe colourimetric assays and the results revealed that AuNPs solution with pCR-pork-mtDNA plasmid and pork DNA remained red, whereas the solution of the rest samples changed to colourless, indicating the aggregation of AuNPs (Fig. 4B). The AuNPs-probe solutions with LAMP products were determined with UV-Vis spectroscopy. The spectroscopic results agreed with agarose gel electrophoresis and visual observation as the unchanged A520/A650 ratios were seen in pCR-pork-mtDNA plasmid and pork DNA samples, but the A520/A650 ratios were decreased in non-pork DNA samples and NTC (Fig. 4C). Taking these data together, the LAMP-AuNP-probe assay developed was highly specific for pork DNA detection.

**Fig. 4.**
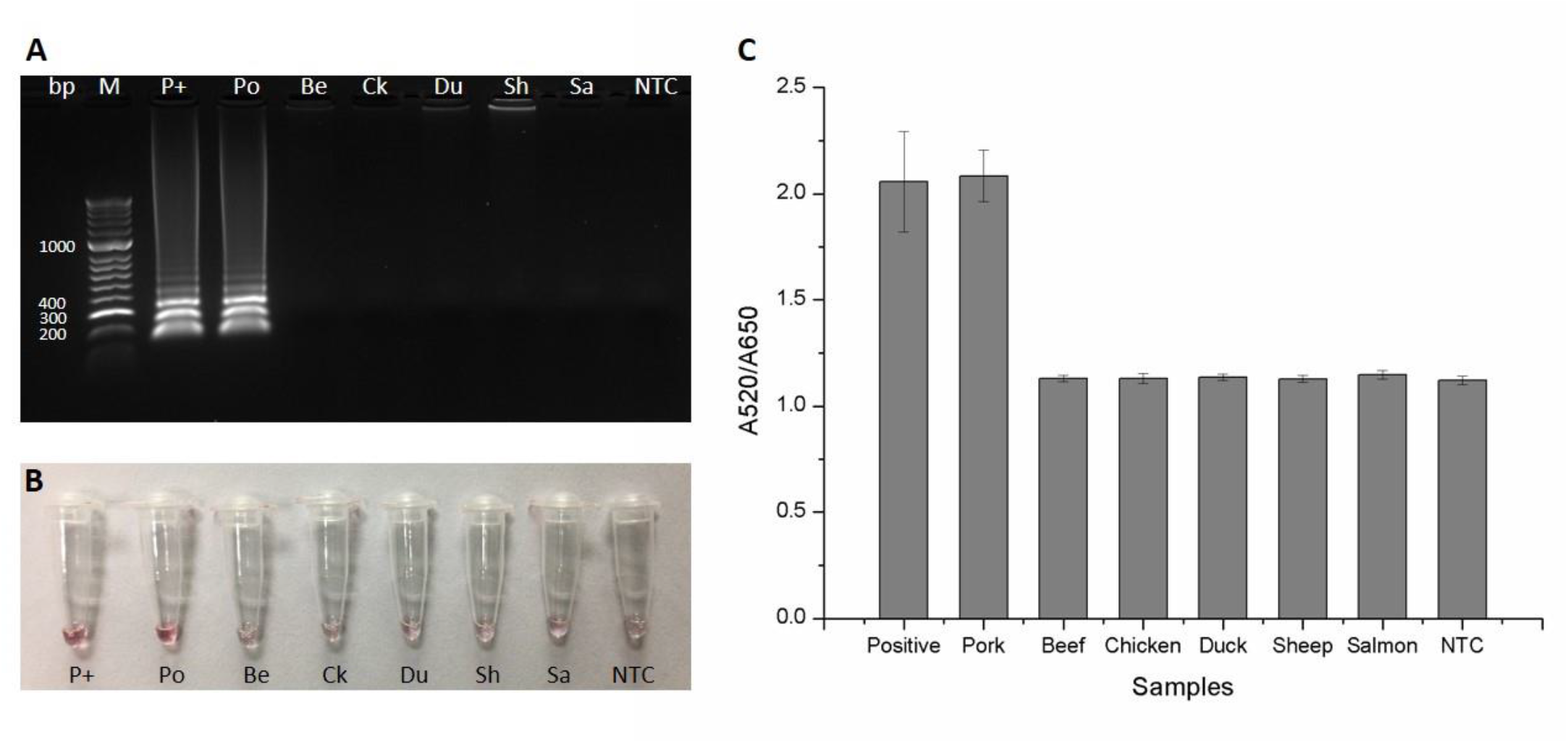
Specificity of LAMP-AuNPs-probe colourimetric assays for genomic DNA from different meat species. (A) LAMP products analysed by 2% agarose gel electrophoresis. (M), HyperLadder 50 bp DNA marker (Bioline, UK); (P+), 25 ng of pCR-pork-mtDNA plasmid as a positive control; (Po), 40 ng of pork DNA; (Be), 40 ng of beef DNA; (Ck), 40 ng of chicken DNA; (Du), 40 ng of duck DNA; (Sh), 40 ng of sheep DNA; (Sa), 40 ng of salmon DNA and (NTC), ddH_2_O as a no-template control. (B) Digital photograph showing AuNPs-probe solution colour with LAMP products amplified from gDNA from different meat species. (C) UV-Vis spectroscopic results (A520/A650) for meat-species specificity test of LAMP-AuNPs-probe assays. Data are means ± SD (n = 8).

Primer-specific PCR and RT-PCR were also conducted to compare the specificity for pork identification with the LAMP-AuNP-probe assay. The specific PCR was performed using a pair of primers specific to pork D-loop mitochondrial DNA, which was obtained from Montiel-Sosa et al. (2000). The primers and probe used for RT-PCR provided by Köppel et al. (2011) were specific for pork beta-actin gene. Both primer-specific PCR and RT-PCR showed high specificity for pork identification as the positive detection was only found in the reaction containing pork DNA (Fig. S6 and S7). The specificity of LAMP-AuNP-probe assay was equivalent to the primer-specific PCR and RT-PCR as the same detection results for pork authentication were observed (Table S1).

### Sensitivity of the LAMP-AuNPs colourimetric assay

Sensitivity of the LAMP-AuNPs colorimetric assay was also studied to determine a limit of detection of the assay, which is the lowest concentration of pork DNA template that can be detected. A range of pork DNA concentrations from 10^2^ to 10^−4^ ng was prepared and utilised to perform LAMP assays. The results from agarose gel electrophoresis revealed that LAMP products were clearly observed with the initial pork DNA template at a concentration of 10^2^ – 10^−1^ ng (Fig. 5A). The products almost disappeared at 10^−2^ ng of pork DNA and were undetectable at a concentration of 10^−3^ – 10^−4^ ng (Fig. 5A). After testing LAMP products with AuNPs colourimetric assays, the data obtained from visual observation (Fig. 5B) and UV-Vis spectroscopy (Fig. 5C) showed that at pork DNA concentrations of 10^2^ – 10^−1^ ng, no AuNPs-probe aggregation was observed and the A520/A650 ratios remained unchanged. However, at pork DNA concentrations between 10^−2^ and 10^−4^ ng, the solution of AuNPs-probe turned colourless (Fig. 5B), indicating the AuNPs aggregation with the decrease of A520/A650 ratios (Fig. 5C). The limit of detection (LOD) for LAMP-AuNPs-probe assays in this work was as low as 0.1 ng (=100 pg) of pork gDNA template. This LOD falls in the nM - pM range, which is acceptable as compared to other published pork identification methods (Table 2).

**Fig. 5.**
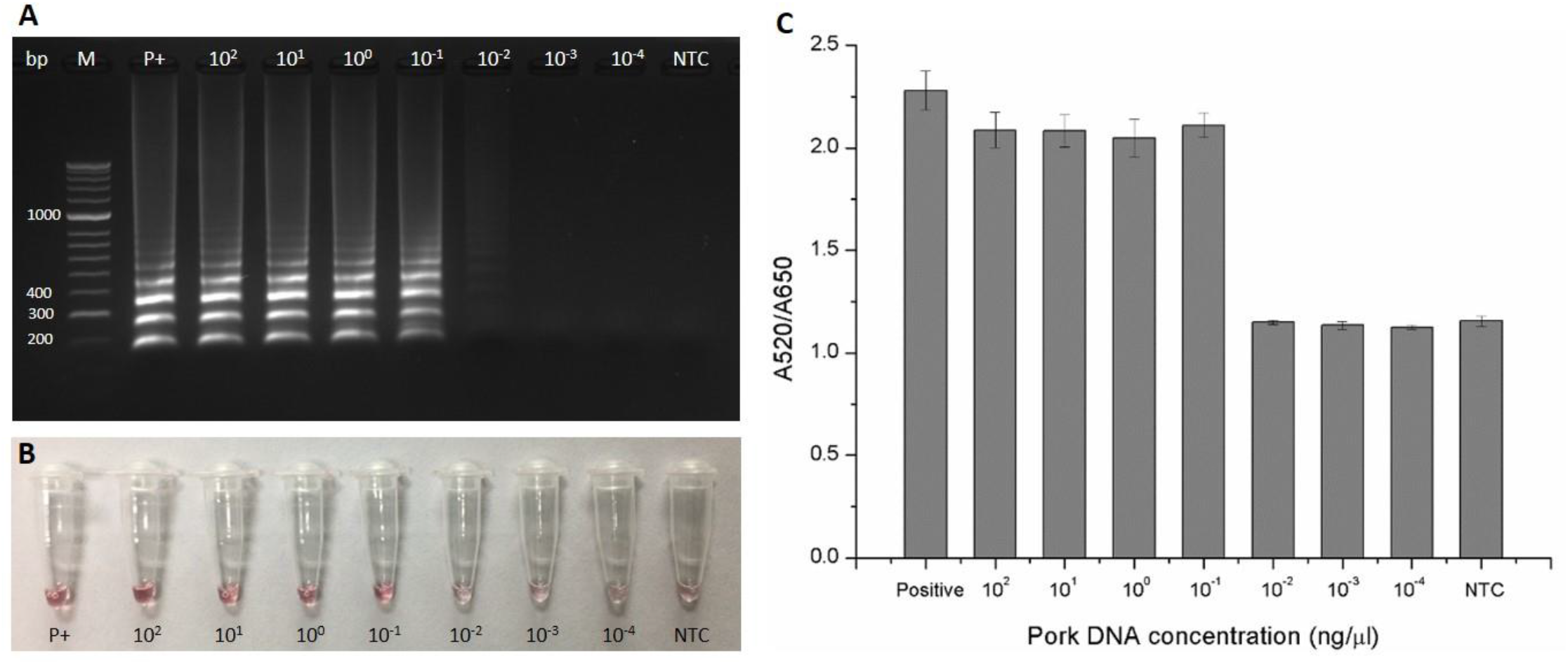
Sensitivity of LAMP-AuNPs-probe colourimetric assays tested with different concentrations of pork genomic DNA. (A) LAMP products analysed using 2% agarose gel electrophoresis. (M), HyperLadder 50 bp DNA marker (Bioline, UK); (P+), 25 ng of pCR-pork-mtDNA plasmid as a positive control; (10^2^ – 10^−4^), pork DNA at the concentration of 10^2^ – 10^−4^ ng and (NTC), ddH_2_O as a no-template control. (B) Digital photograph showing AuNPs-probe solution colour with LAMP products amplified from different concentrations of pork gDNA (C) UV-Vis spectroscopic results (A520/A650) for sensitivity test of LAMP-AuNPs-probe assays. Data are means ± SD (n = 6).

**Table 2.** Sensitivity of PCR-based assays for pork DNA detection.

| **Methods** | **Target genes** | **Limit of detection for pork DNA** | **References** |
| --- | --- | --- | --- |
| Specific PCR | Mitochondrial D-loop gene | 1 pg/μl | Man et al. (2012) |
| Specific PCR | Mitochondrial D-loop gene | 10 pg | Karabasanavar et al. (2014) |
| Multiplex PCR | Species-specific nuclear DNA (nDNA) | 500 pg | Wang et al. (2020) |
| RT-PCR | Mitochondrial cytochrome-b gene | 1 pg | Orbayinah et al. (2019) |
| RT-PCR | Pig mitochondrial DNA | 100 pg | Farrokhi and Joozani (2011) |
| RT-PCR | 12S rRNA gene | 10 pg | Rodríguez et al. (2005) |
| RT-PCR | Mitochondrial D-loop DNA | 0.1 pg | Kim et al. (2016) |
| Multiplex RT-PCR | Beta-actin gene | 150 pg | Cheng et al. (2014) |
| Molecular Beacon Probe RT-PCR | Cytochrome b gene | 0.1 pg | Yusop et al. (2012) |
| ddPCR | ATPase gene | 2.2 pg | Shehata et al. (2017) |
| Duplex ddPCR | Beta-actin gene | 100 pg/μl | Cai et al. (2017) |
| RT-LAMP | Mitochondrial D-loop gene | 1 pg | Lee et al. (2016) |
| RT-LAMP | Cytochrome b gene | 1 pg | Yang et al. (2014) |
| Gold nanoparticle sensor | Cytochrome b gene | 400 pg/μl | Ali et al. (2012) |
| LAMP-AuNP-oligoprobe assay | Mitochondrial DNA | 100 pg | This work |

### Challenge of the LAMP-AuNPs-probe assay with processed food samples

The LAMP-AuNPs-probe assays were demonstrated for their capability of detecting pork content in process-food products collected from supermarkets in Pathum Thani, Thailand. All the processed food products were tested for the presence of pork content by droplet digital PCR (ddPCR) (Temisak et al. 2020) prior to LAMP-AuNPs-probe colourimetric assays (Table S2). In Fig. 6A, LAMP assays with agarose gel electrophoresis showed that the amplified products were detected in a sample of pCR-pork-mtDNA plasmid, two samples of pork sausage and a sample of pork ball. However, no LAMP products were seen in non-pork products which were chicken burger, beef burger and shrimp dumpling (Fig. 6A). The data from AuNPs-probe colourimetric assays also agreed with the results of agarose gel electrophoresis. As seen in Fig. 6B, the AuNPs-probe remained red for the samples containing pork DNA (pCR-pork-mtDNA plasmid, two pork sausage samples and pork ball), indicating no aggregation of AuNPs. These pork DNA containing samples also showed higher ratios of A520/A650 in UV-Vis spectroscopic measurement (Fig. 6C). On the other hand, non-pork DNA containing samples (chicken burger, beef burger, shrimp dumpling and NTC) showed the change of AuNPs colour to colourless (Fig. 6B) and lower ratios of A520/A650 (Fig. 6C) as anticipated, indicating the AuNP aggregation. This means that the LAMP-AuNPs-probe assays developed in this study are capable of detecting pork content in commercial processed-food products in the markets.

**Fig. 6.**
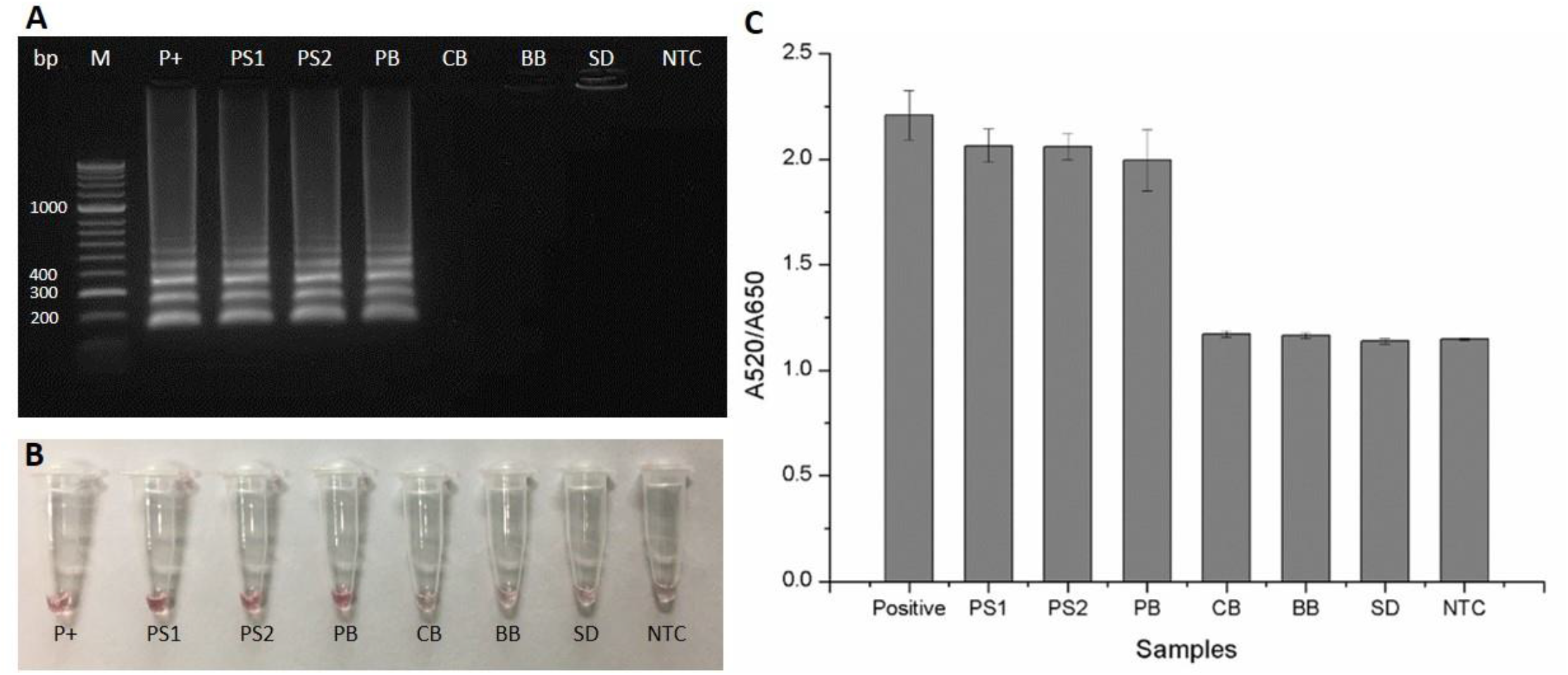
The challenge of LAMP-AuNPs-probe colourimetric assay with processed-food products. (A) LAMP products analysed by 2% agarose gel electrophoresis. (M), HyperLadder 50 bp DNA marker (Bioline, UK); (P+), 25 ng of pCR-pork-mtDNA plasmid as a positive control; 10 ng of DNA extracted from (PS1), pork sausage no.1; (PS2), pork sausage no.2; (PB), pork ball; (CB), chicken burger; (BB), beef burger; (SD), shrimp dumpling and (NTC), ddH_2_O as a no-template control. (B) Digital photograph showing AuNPs-probe solution colour with LAMP products amplified from processed-food product DNA (C) UV-Vis spectroscopic results (A520/A650) for LAMP-AuNPs-probe assays tested with processed food samples. Data are means ± SD (n = 6).

### Efficiency and limitations of the assay

In this work, a new pork authentication method based on a combination of loop-mediated isothermal amplification and AuNP-nanoprobe colourimetric assay has been successfully developed. The technique has been proven for its fast processing time with 45 min for LAMP assay (Fig. S3) plus 45 min for AuNP-oligoprobe assay. The LAMP-AuNP-nanoprobe assay also offers high specificity to the pork content and exhibits no cross reactivity with other meat species (Fig. 4 and 6). The assay also shows high sensitivity with the limit of detection of pork genomic DNA as low as 100 pg (Fig. 5).

Conventional PCR methods such as RAPD-PCR, RFLP-PCR and PCR with sequence-specific primers have been reported for pork identification (Ballin et al. 2009; Man et al. 2012; Mane et al. 2013). Although proven to be specific and sensitive to the targets, the PCR based techniques are cumbersome due to the requirements of special equipment such as thermal cyclers. They may also require an additional step after performing PCR like gel electrophoresis in order to reveal the outcome (Ali et al. 2012; Calvo et al. 2001a; Calvo et al. 2001b; Man et al. 2012; Mane et al. 2013; Murugaiah et al. 2009). The use of LAMP together with AuNP-nanoprobe method developed in this study could be a promising alternative as it can be conducted at a constant temperature and the detection can be clearly observed by the naked eyes. The LAMP-AuNP-probe assay, however, has its limitations as it delivers qualitative information and is less sensitive when compared with the quantitative PCR techniques (Table 2). Real time PCR (RT-PCR) and digital PCR (dPCR) are the two PCR-based methods that can provide quantitative data with high specificity and sensitivity (Ballin et al. 2009; Cai et al. 2017; Farrokhi and Joozani 2011; Floren et al. 2015; Orbayinah et al. 2019; Rodríguez et al. 2005; Shehata et al. 2017). Nevertheless, both RT-PCR and dPCR require relatively expensive reagents as well as special devices to perform the PCR reactions, which may not be affordable for laboratories with limited budgets (Ali et al. 2012; Basu 2017). The LAMP-AuNP-probe assay could thus be a good solution for this problem as it is inexpensive and requires only a simple device that can control the temperature such as a water bath or a heating block.

## Conclusion

In the present study, a new analytical method using a loop-mediated isothermal amplification combined with AuNP-oligoprobe colourimetric assay has been developed successfully for meat authentication, especially to distinguish pork from other meat products. The assay was highly specific to pork DNA content without cross reactivity with other meat species. It is highly sensitive with a limit of detection as low as 100 pg of pork DNA. In addition, this method has been proven to be rapid, easy-to-use and cost-effective, as it only requires simple temperature-controllable devices for the experimentation. No special detection system is required as the transition of colloidal AuNPs colour can be readily visually observed. Hence, it is anticipated that the developed method could become a viable alternative for laboratories with limited budget and equipment, rendering authentication of pork or halal food more convenient, yet still precise and reliable.

## Acknowledgements

This work was partially supported by the Global Partnership Research Grant by the Thailand Science Research and Innovation (TSRI). N.S. was also supported by the Materials Science Research Center (MSRC), Chiang Mai University, and the National Research Council of Thailand (Grant: 2556NRCT51390, 2558NRCT350269, and NRCT-PARB/39/2561).

## Compliance with ethical standards

### Conflict of interest

Pattanapong Thangsunan declares that he has no conflict of interest. Sasithon Temisak declares that she has no conflict of interest. Phattaraporn Morris declares that she has no conflict of interest. Leonardo Rios-Solis declares that he has no conflict of interest. Nuttee Suree declares that he has no conflict of interest.

### Ethical approval

This article does not contain any studies with human participants or animals performed by any of the authors.

### Informed consent

Not applicable.

## Notes

### Competing Interest Statement

The authors have declared no competing interest.

